# Cell surface architecture of the cultivated DPANN archaeon *Nanobdella aerobiophila*

**DOI:** 10.1101/2023.10.27.564410

**Authors:** Shingo Kato, Yuhei O. Tahara, Yuki Nishimura, Katsuyuki Uemastu, Takahiro Arai, Daisuke Nakane, Ayaka Ihara, Takayuki Nishizaka, Wataru Iwasaki, Takashi Itoh, Makoto Miyata, Moriya Ohkuma

## Abstract

The DPANN archaeal clade includes obligately ectosymbiotic species. Their cell surfaces potentially play an important role in the symbiotic interaction between the ectosymbionts and their hosts. However, little is known about the mechanism of the ectosymbiosis. Here, we show cell surface structures of the cultivated DPANN archaeon *Nanobdella aerobiophila* strain MJ1^T^ and its host *Metallosphaera sedula* strain MJ1HA, using a variety of electron microscopy techniques, i.e., negative-staining transmission electron microscopy (TEM), quick-freeze deep-etch (QFDE) TEM, and 3D electron tomography. The thickness, unit size, and lattice symmetry of the S-layer of strain MJ1^T^ were different from those of the host archaeon strain MJ1HA. Genomic and transcriptomic analyses highlighted the most highly expressed MJ1^T^ gene for a putative S-layer protein with multiple glycosylation sites and immunoglobulin-like folds, which has no sequence homology to known S-layer proteins. In addition, genes for putative pectin lyase- or lectin-like extracellular proteins, which are potentially involved in symbiotic interaction, were found in the MJ1T genome based on in silico 3D protein structure prediction. Live cell imaging at the optimum growth temperature of 65°C indicated that cell complexes of strains MJ1^T^ and MJ1HA were motile, but sole MJ1^T^ cells were not. Taken together, we propose a model of the symbiotic interaction and cell cycle of *Nanobdella aerobiophila*.

**Importance:** DPANN archaea are widely distributed in a variety of natural and artificial environments, and may play a considerable role in the microbial ecosystem. All of the cultivated DPANN archaea so far need host organisms for their growth, i.e., obligately ectosymbiotic. However, the mechanism of the ectosymbiosis by DPANN archaea is largely unknown. To this end, we performed a comprehensive analysis of the cultivated DPANN archaeon, *Nanobdella aerobiophila*, using electron microscopy, live cell imaging, transcriptomics, and genomics including 3D protein structure prediction. Based on the results, we propose a reasonable model of the symbiotic interaction and cell cycle of *Nanobdella aerobiophila*, which will enhance our understanding of the enigmatic physiology and ecological significance of DPANN archaea.

## Introduction

The DPANN archaeal clade has been proposed as a superphylum-level archaeal clade in 2013 (1). Although DPANN archaea are widely distributed on Earth, little is known about their ecological roles because most of them are yet uncultivated (2, 3). Culture-independent metagenomics/single cell genomics suggest most of the DPANN archaea represent symbiotic lifestyles (2, 3). Several cultivates of DPANN archaea have been reported so far, i.e., ‘*Nanoarchaeum equitans’*, *Nanobdella aerobiophila,* and some others in the phylum *Nanobdellota* (formerly called*‘Nanoarchaeota’*)(4–9), and several species in the phylum-level clades *‘Microcaldota’/‘Micrarchaeota’* (10, 11) and *‘Nanohaloarchaeota’* (12, 13). These cultivated members are commonly small in cell and genome sizes, and obligate ectosymbionts requiring host organisms for their growth. Cells of DPANN archaea directly attach to the cell surface of their hosts. Their small genomes lack the synthesis pathway of primary materials, such as nucleotides, lipids, and even ATP in some cases. Thus, the DPANN archaea probably obtain these primary materials from their hosts. In the case of ‘*Nanoarchaeum equitans’* and its host *Ignicoccus hospitalis*, host-specific proteins have been detected in the cell interior of its symbiont (14). A hole in the cytoplasmic membranes connected between the cytoplasmic areas of a symbiont and a host has been observed in the complex of ‘*Nanoarchaeum equitans’–I. hospitalis* cells and also of *Nanobdella aerobiophila*–*Metallosphaera sedula* cells (5, 14), which is likely to the passage for material transfer. However, it is largely unclear how the DPANN archaeal symbionts recognize, attach to, and pierce their hosts.

Cell surfaces potentially play an important role in the construction of ectosymbiotic relationships between DPANN archaea and their hosts. The cell surface of most archaea including DPANN archaea is covered by an S-layer (15, 16) that generally consists of the most abundantly expressed proteins in each archaeon (17). The archaeal S-layer exhibits either oblique (p1 and p2), square (p4), or hexagonal (p3 and p6) lattice symmetry (16). In particular, the S-layers of ‘*Nanoarchaeum equitans’* and *‘Micrarchaeum harzensis’* in the DPANN clade have been reported to represent a hexagonal lattice (p3/p6) (4, 18, 19). Putative genes encoding S-layer proteins have been identified in the genomes of ‘*Nanoarchaeum equitans’* (NEQ300) (20) and *‘Micrarchaeum harzensis’* (Micr_00292) (19). Indeed, these genes are highly expressed in the transcriptomes and proteomes of the two DPANN archaea (10, 21, 22). In addition to S-layers, DPANN archaea often represent archaellum-like cell surface filaments. Archaella are involved in motility, surface adhesion, cell–cell interaction, and biofilm formation (23, 24), and belong to the type-IV filament superfamily including bacterial type-IV pili (24). Moreover, other extracellular filaments, e.g., threads and bundling pili, have been found in archaea (25, 26). The S-layers and filaments are generally highly glycosylated and represent globular folds rich in β-strands (e.g., immunoglobulin (Ig) or polycystic kidney disease (PKD) fold), of which feature may be involved in protein-protein and protein-ligand interaction (27). Indeed, genes for the cell surface proteins with such features have been found in DPANN genomes, and predicted to be involved in surface interactions (2, 3, 12, 13, 28).

Previously, we have reported the cultivation of a DPANN archaeon, *Nanobdella aerobiophila* strain MJ1^T^, which can grow under aerobic conditions with *Metallosphaera sedula* strain MJ1HA as its host (5). The presence of S-layers and filaments on the cell surface of the strains MJ1 and MJ1HA has been revealed by transmission electron microscopy (TEM) (5). Here we report more details of the cell surfaces of these strains using quick-freeze deep-etch (QFDE)-TEM and 3D electron tomography, in addition to genomic and transcriptomic analyses. Our results will provide insights into the symbiotic mechanism of DPANN archaea.

## Materials and Methods

### Strains and culture conditions

Coculture of *Nanobdella aerobiophila* strain MJ1^T^ (= JCM 33616^T^) and *Metallosphaera sedula* strain MJ1HA (=JCM 33617) and pure-culture of strain MJ1HA were obtained from the Japan Collection of Microorganisms (JCM), and maintained by subcultivation at pH 2.5 and 65°C under the aerobic conditions as described previously (5).

### Motility observation

The cell motility of strains MJ1^T^ and MJ1HA was observed at 65°C using a live-imaging system. The cell culture was poured into a tunnel chamber assembled by taping a coverslip, and both ends of the chamber were sealed with nail polish to keep from drying the sample, as previously described (29). Cell behavior was visualized under an inverted microscope (IX71, Olympus) equipped with an objective lens (LUCPLFLN 40×PH, Olympus), a CMOS camera (DMK33U174, Imaging Source, Germany), and an optical table (HAX-0806, JVI, Japan). The microscope stage was heated with a thermoplate (TP-110R-100, Tokai Hit, Japan). Projections of the images were captured as greyscale images with the camera under 0.05-s resolution and converted into a sequential TIF file without any compression. All data were analyzed by ImageJ version 1.48 (30) and its plugins, TrackMate, Particle Tracker, and Multitracker.

### Scanning electron microscopy and ultra-thin section transmission electron microscopy

The images of ultra-thin section transmission electron microscopy (TEM) and of scanning electron microscopy (SEM) were obtained in the previous study (5). In brief, for SEM, the collected cells were fixed using a rapid freezing/freeze-substitution method, and were coated with a thin layer (30nm) of osmium. The coated samples were observed using field-emission scanning electron microscopy (FE-SEM). For ultra-thin section TEM, the fixed cells in the polymerized resins were ultra-thin sectioned at a thickness of 70 nm using an ultra-microtome, strained with 2% uranyl acetate, and observed by a transmission electron microscope. All of the images used in this paper were not shown in the previous paper.

### Negative-staining transmission electron microscopy

Although some images of negative-staining TEM of MJ1^T^ and MJ1HA cells have been reported previously (5), in this study, we performed the negative-staining TEM again to obtain additional information for cell morphology. Cells of MJ1^T^ and MJ1HA were collected from a 3-day coculture by centrifugation at 15,000 × g for 4 min. The pellet was suspended in buffer containing 10 mM (NH_4_)_2_SO_4_, 2 mM KH_2_PO_4_, 2 mM MgSO_4_, and 0.5 mM CaCl_2_ adjusted pH to 2.5 with 10 N H_2_SO_4_ at one-tenth of the original culture volume. Sample preparation for negative-staining EM followed the same protocol described previously (31). Carbon-coated EM grids were glow-discharged by a PIB-10 hydrophilic treatment device (Vacuum Device, Japan). The cell suspension was placed on the EM grid and kept for 10 min at room temperature (RT), and then chemically fixed by 0.5% (w/v) glutaraldehyde in the buffer for 10 min at RT. After washing three times with the buffer, the grid was stained with 2% ammonium molybdate (w/v). Samples were observed under a transmission electron microscope (JEM-1400, JEOL, Japan) at 100 kV. A charge-coupled device camera captured EM images.

### Quick-freeze deep-etch electron microscopy

To obtain fine images of the surface structure of the cells, QFDE-TEM was performed as described previously (32). Aliquots of 4-day cocultures of strain MJ1 and MJ1HA and those of 4-day pure culture of strain MJ1HA were used for the QFDE-EM. Cells in the cultures were harvested by centrifugation at 10,000 x g for 5 min. In brief, the cell suspension was frozen by a CryoPress (Valiant Instruments, St. Louis, MO), fractured and etched at -104°C in a freeze-etching device (JFDV; JEOL Ltd., Akishima, Japan), and rotary-shadowed by platinum. The replicas were floated off, rinsed in ultrapure water, cleaned with a bleach, and observed by a transmission electron microscope (JEM-1010; JEOL). The corresponding power spectra of selected areas were produced through Tukey Window, Fast Fourier Transform (FFT), and contrast enhancement. The spectra were converted by inverse FFT to images emphasized for periodic structure.

### 3D electron tomography

To analyze the 3D structure of the MJ1^T^ and MJ1HA cells, two methods were used as follows: (i) Scanning transmission electron microscopy (STEM) tomography method was used to obtain a fine 3D model of the boundary between MJ1^T^ and MJ1HA cells; (ii) Serial cross-sectional images were obtained by focused ion beam (FIB)-SEM to reconstruction a model of the whole structure of the MJ1–MJ1HA cell complexes. Cells of MJ1^T^ and MJ1HA were collected from a 9-day coculture by centrifugation at 10,000 × g for 5 min. After removing the supernatant, the centrifuged cells were added to 4% agarose LGT (low gelling temperature)(Nacalai Tesque Inc., Kyoto, Japan) diluted in medium, then suspended and applied to a flat-specimen carrier and frozen in a high-pressure freezing device (EM-PACT; Leica). The frozen samples were substituted with 2% OsO_4_ in acetone for 3 days at –80°C. To facilitate the extraction of intracellular microstructures during tomogram preparation, en bloc staining was performed using methanol for 1 h at 4°C, and then the stained samples were embedded in epoxy resin (33).

For STEM tomography, the semi-thin sections (thickness: 500 nm) were cut from the embedded samples by a diamond knife (Histo; Diatome, Nidau, Switzerland), and collected on formvar-coated copper grids. The grids were stained with a 2% uranyl acetate solution for 20 min and a lead stain solution for 10 min, and then coated with carbon (about 15 nm thickness). Tomography data was acquired in a tilted series with a TEM (Tecnai 20; ThermoFisher Scientific, Waltham, USA). The detailed conditions were modified from the previous report (34). In brief, acceleration voltage: 200 kV, acquisition tilt angle: ±70°, acquisition image step: θ cosθ. STEM images were taken with an annual dark-field detector (camera length: 285 mm). In addition, to reduce the area of missing tomography data in the Z direction, dual axial tomography (35) was performed by acquiring a single tilt image and then rotating the sample 90° to acquire the same area. Then, each series of tilted images was integrated into a single serial tomogram using Inspected 3D (ThermoFisher Scientific). FIB/SEM imaging data was acquired by a FIB/SEM (Helios G4 UX: ThermoFisher Scientific) to obtain a series of serial cross-sections. The detailed conditions were modified from the previously described method (36). In brief, after milling with a gallium ion beam at an acceleration voltage of 30 kV, current of 0.26 nA, and 15 nm/step, images were acquired with a mirror detector (backscattered electron image) at an acceleration voltage of 2 kV, current of 1.6 nA. These processes were repeated to obtain a serial cross-sectional image series.

Both single serial tomograms obtained by STEM tomography and FIB/SEM imaging were processed using Amira v6.3.0 (ThermoFisher Scientific). The final tomographic model was obtained as previously described (37).

### Genome and transcriptome analysis

The complete genome sequences of *Nanobdella aerobiophila* strain MJ1^T^ (NCBI accession number, AP019769) and *Metallosphaera sedula* strain MJ1HA (AP019770) were analyzed in this study. Protein domains for all protein-coding regions (CDSs) in the genomes were predicted using InterProScan version 5.59-91.0 (38), including TMHMM version 2.0 (39) and Phobius version 1.01 (40) for transmembrane (TM) and signal peptide sequences. In addition, SignalP 6.0 (41) was also used for signal peptide sequence prediction. 3D protein structures were predicted on the ColabFold website (version 1.5.2)(42) or by a stand-alone AlphaFold version 2.2.0 (43). Using the PDB formatted files, the 3D structures of proteins were viewed in ChimeraX version 1.6.1 (44). Homology searching against the NR database was performed by BLASTp on the NCBI website. N- and O-glycosylation sites are predicted by GlycoPP version 1.0 (45). For the above analysis, default parameters were used except where otherwise noted.

Cells of the MJ1–MJ1HA coculture and MJ1HA-pure culture from 9-day cultures were harvested as described above, and used for the following RNA-seq transcriptome analysis. The harvested cell samples were treated in cell lysis buffer (100 mM Tris-HCl (pH 7.5), 100 mM NaCl, 0.5 mM EDTA, 0.07% SDS, and proteinase K) at 40°C for 30 min, and total RNAs were extracted from the samples using a NucleoSpin RNA kit (Macherey-Nagel, Düren, Germany). cDNAs were synthesized from the extracted RNAs using a PrimeScript Double Strand cDNA Synthesis Kit (Takara Bio, Kusatsu, Japan) with 9 mer random primers. The library construction and MiSeq sequencing were performed as described previously (5). Counting reads for CDSs were performed using featureCounts version 2.0.1 (46), and values of transcripts per million (TPM) were calculated as reported previously (47).

## Results and discussion

### Electron Microscopy

As previously reported (5), the ultra-thin section TEM indicated the presence of S-layers of both MJ1^T^ and MJ1HA cells (Fig. 1A and B; Fig. S1). The S-layer of the host archaeon *Metallosphaera sedula* strain MJ1HA represented regular protrusions, whereas that of *Nanobdella aerobiophila* MJ1^T^ looked smoother. The difference in the appearance of the S-layer between the strains MJ1^T^ and MJ1HA was also clearly shown in QFDE-TEM images of replicas of the cell surfaces (Fig. 1C; Fig. S2).

**Fig. 1.**
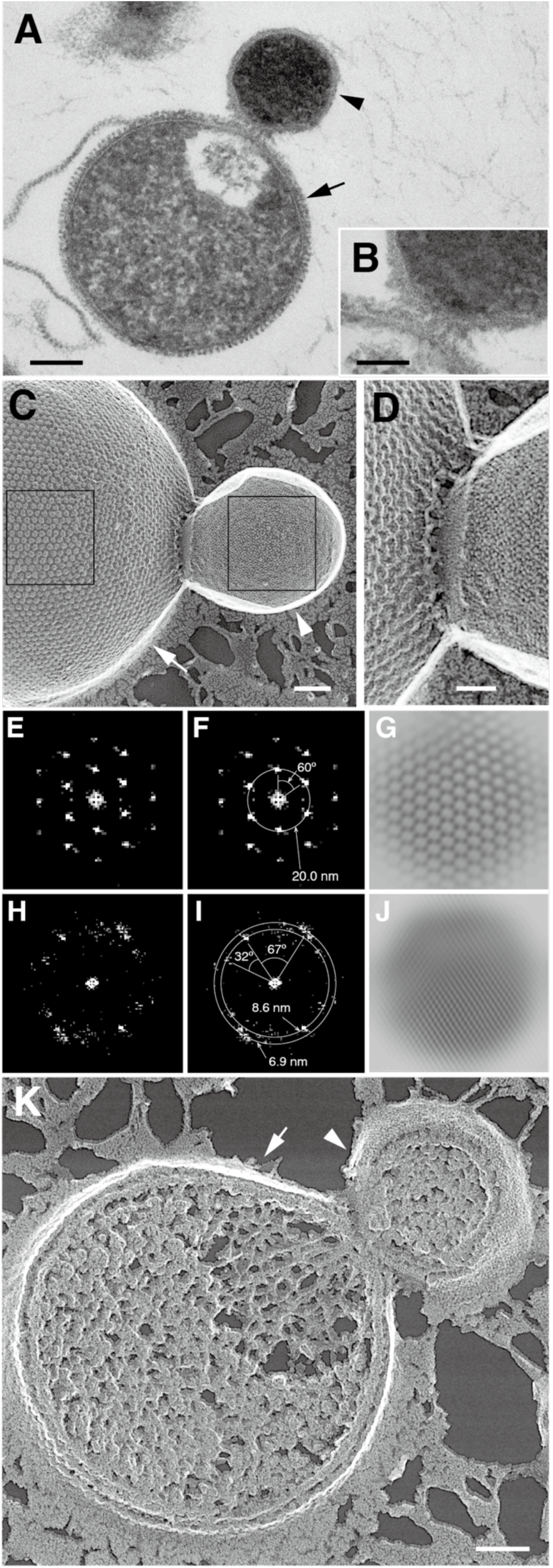
Electron micrograph of interior and surface of *Nanobdella aerobiophila* cells. **(A)** A TEM image of an ultra-thin section of a cell complex of *Nanobdella aerobiophila* strain MJ1^T^ (arrowhead) and its host archaeon *Metallosphaera sedula* strain MJ1HA (arrow), and **(B)** the enlarged view of the contact area between the cells. **(C)** A QFDE-TEM image of a replica of cell surfaces of strains MJ1^T^ (arrowhead) and MJ1HA (arrow), and **(D)** the enlarged view of the contact area between the cells. Corresponding FFT power spectra of cell surfaces of strains MJ1HA **(E and F)** and MJ1^T^ **(H and I)** from the boxes (left for strain MJ1HA, right for strain MJ1^T^) of the panel C. **(G and J)** Inverse FFT image of extracted spots from the panels E and H. **(K)** A cross-fracture QFDE-TEM image of a cell complex of strains MJ1^T^ (arrowhead) and MJ1HA (arrow). Bars, 200 nm (A), 100 nm (B, C and K), and 50 nm (D).

For the MJ1HA cell surface, each of the S-layer protein units was hexagonal with 18–20 nm in a flat-to-flat distance (Fig. 1C). The MJ1HA S-layer exhibited hexagonal lattice symmetry with 19–20 nm of a center-to-center distance revealed by the FFT analysis (Fig. 1E–G; Fig. S3). There were ∼5 nm spaces between each unit. No clear difference in the above features between the MJ1HA cells in coculture and pure culture was observed (Fig. 1C; Fig. S3). It is unclear if the hexagonal lattice symmetry of MJ1HA represents p3 or p6 at present, although those of the type strain TH2^T^ of *Metallosphaera sedula* and others in the order *Sulfolobales* have been reported to be p3 (16).

In contrast, the S-layer units of strain MJ1^T^ were irregular in shape, and smaller in size (3–5 nm in the major axis; over 10 nm in some cases)(Fig. 1C). There were ∼2 nm spaces between each unit. The FFT analysis showed an oblique pattern of the MJ1 S-layer lattice symmetry (possibly p1/p2) with an 7–9 nm of center-to-center distance (Fig. 1H–J; Fig. S3), which seems to be different from the hexagonal lattice (p3/p6) of ‘*Nanoarchaeum equitans’* and *‘Micrarchaeum harzensis’* (4, 18, 19).

Notably, the cell surface of MJ1^T^ at the contact sites with MJ1HA cells was relatively flat, as compared with other whole S-layer areas (Fig. 1D; Fig. S2). The S-layer units on the flat areas could not be confirmed because the unit size might be much smaller. In some cases, filamentous structures (∼15 nm in width) were observed at the contact sites (Fig. S2), which was also observed by the ultra-thin section TEM (Fig. 1B; Fig. S1E). Such filamentous structures have been also reported in the contact sites between other DPANN archaea and their hosts (10, 11).

Cross-fracture QFDE-TEM images showed the cytoplasmic voids at the part of MJ1HA cells next to the attached MJ1 cells (Fig. 1K; Fig. S4). This is consistent with the previous report showing the low electron density at such cytoplasmic areas of MJ1HA cells, which have been interpreted to be the result of sucking by MJ1^T^ (5).

The thickness of the S-layer of MJ1^T^ cells (15–20 nm) was generally thinner than that of MJ1HA cells (25–35 nm), as shown by both QFDE-TEM and ultra-thin section TEM (Fig. 1; Fig. S1 and S4). The thickness of the cell membrane was 3–4 nm for both strains.

The STEM tomography confirmed the presence of holes penetrating the cell membranes of MJ1^T^ and MJ1HA (Fig. 2A and 2B; Movie S1 and S2), as previously reported by ultra-thin section TEM (5)(Fig. S1I). The holes were round and ∼30 nm in diameter, implying that smaller proteins (even ∼25 nm-size ribosomes (48, 49)) could be passed through the holes. At the contact sites, a hill-like protrusion of a MJ1 cell was observed both in the STEM and FIB/SEM tomography (Fig. 2A–C; Movie S1–3). The whole image of an MJ1–MJ1HA cell complex was shown by FIB/SEM tomography (Fig. 2C; Movie S3), suggesting that MJ1^T^ cells were attached from all directions to the MJ1HA surface. Two or more cells of MJ1^T^ were tightly adjacent to each other on the host cell surface, indicating that these MJ1^T^ cells were proliferated there. Such proliferated cells were also observed by SEM (Fig. 2D; Fig. S5).

**Fig. 2.**
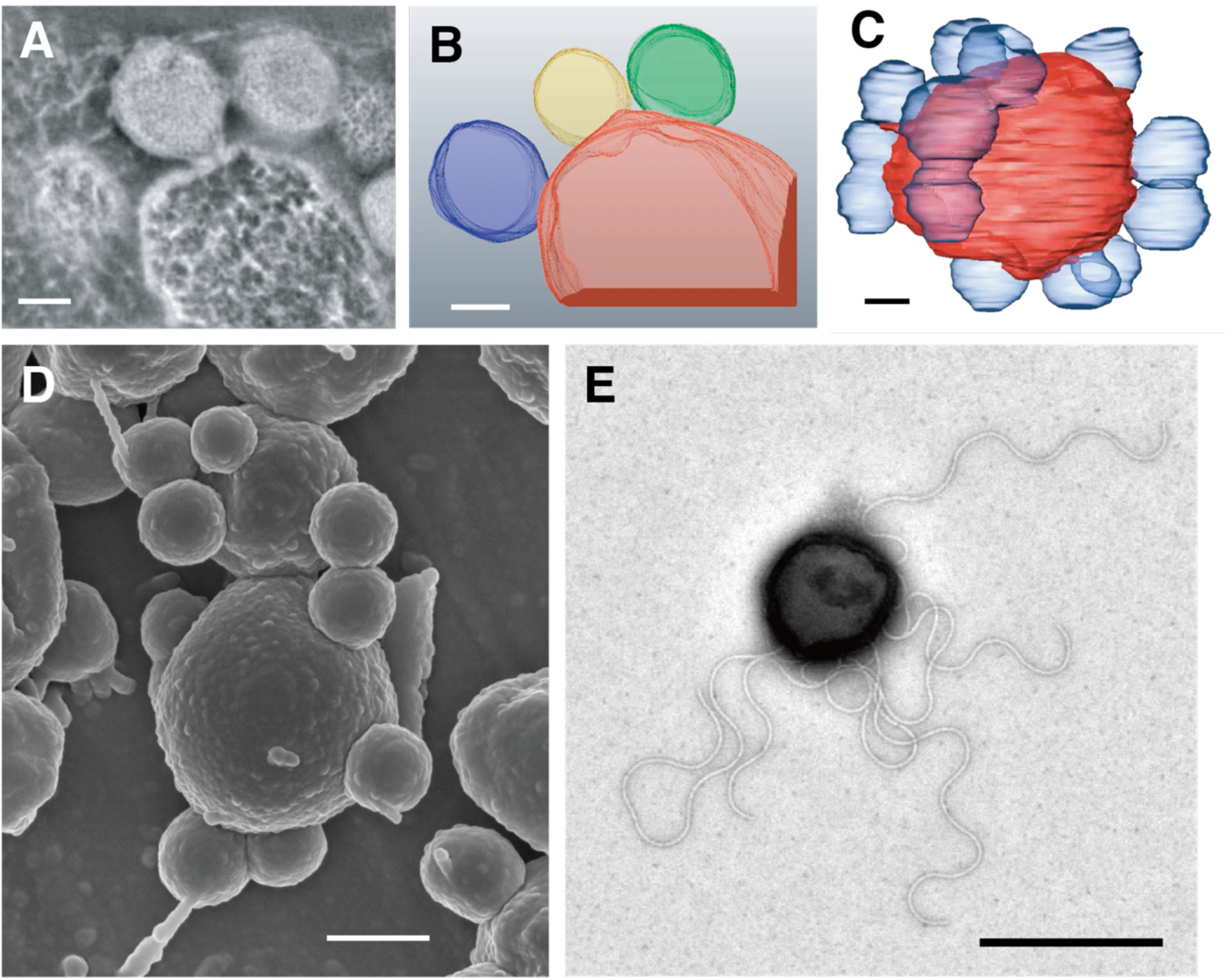
Whole cell complex of *Nanobdella aerobiophila* and its host *Metallosphaera sedula*. **(A)** A slice image of sequential sections of a cell-cell connecting area obtained by STEM tomography. The compressed view of the full slices was provided in Movie S1. **(B)** 3D reconstruction from the full slices by STEM tomography. The full reconstruction was provided in Movie S2. **(C)** A representative image of 3D reconstruction of a whole cell complex of strain MJ1^T^ and MJ1HA obtained by FIB/SEM tomography. The full reconstruction was provided in Movie S3. **(D)** A SEM image of a whole cell complex of strain MJ1^T^ and MJ1HA. Filamentous structures on some MJ1^T^ cells were observed. **(E)** A negative staining TEM image of a cell of strain MJ1^T^. Archaellum-like filaments were observed around the cell. Bar, 200 nm (A, B, C), 500 nm (D, E).

Our previous negative-staining TEM has indicated the presence of archaellum-like filamentous structures (12–14 nm in width) of strain MJ1HA (5). In this study, the newly performed negative-staining TEM indicated that such archaellum-like structures were also present in MJ1^T^ (Fig. 2E; Fig. S6). The MJ1^T^ archaellum-like filaments were 7–10 nm in width, and over 1 µm in length in some cases. It should be noted that we have previously shown the presence of extracellular filamentous structures of MJ1^T^ (50-80 nm in width) by SEM (Fig. 2D)(5), while it is still unclear if these filaments observed in the SEM and negative-staining TEM were identical to each other.

### Transcriptomic analysis and protein structure prediction

We performed the genome and transcriptome analyses to get more information on the cell surface proteins. The complete genome sequences of *Nanobdella aerobiophila* MJ1^T^ and its host archaeon *Metallosphaera sedula* strain MJ1HA have been already determined, and briefly described (5). In this study, we focus on genes for putative cell surface proteins that may play important roles in symbiotic interaction. The genes encoding putative cell surface proteins, including S-layer and archaellum component proteins, found in the MJ1^T^ and MJ1HA genomes are listed in Table 1, as described below in detail. All genes for the MJ1^T^ and MJ1HA genomes with mRNA expression levels are listed in Table S1 and S2. For the MJ1^T^ transcriptome, 735 of the 736 CDSs were expressed in the coculture (Table S1). For the MJ1HA transcriptome, 2560 and 2562 of the 2574 CDSs were expressed in the pure culture and coculture, respectively (Table S2). Regarding protein export to the cell surface, the MJ1HA genome possesses genes related to the Tat system and Sec system (50), but the MJ1^T^ genome possesses genes only for the Sec system. Genes involved in N-glycosylation, e.g., *aglBDE* (51), were found both in the MJ1^T^ and MJ1HA genomes.

**Table 1.** List of the genes encoding putative cell surface proteins, including S-layer and archaellum proteins, found in the MJ1^T^ and MJ1HA genomes. The transcriptomic result was included (*the TPM values and rank in coculture (left) and pure culture (right)).

| Locus_tag | Length (a.a.) | Annotation | #N-glycosylation sites | #Ig-like domains | TM region (start - end a.a.) | Extracellular region (start - end a.a.) | TPM (Rank)* |
| --- | --- | --- | --- | --- | --- | --- | --- |
| **MJ1HA** |  |  |  |  |  |  |  |
| MJ1HA_2063 | 1361 | S-layer protein SlaA | 44 | 9 | 7-29 | 23-1361 | 42447 (1) / 20666 (2) |
| MJ1HA_1540 | 302 | Flagellin ArlB2 | 5 | 1 | 20-42 | 51-302 | 19811 (2) / 16354 (4) |
| MJ1HA_2064 | 416 | S-layer protein SlaB | 8 | 3 | 5-27, 390-412 | 29-391 | 12530 (5) / 4588 (20) |
| **MJ1** |  |  |  |  |  |  |  |
| MJ1_0160 | 1000 | Hypothetical protein | 15 | 5 | 21-43 | 39-1000 | 84646 (1) |
| MJ1_0453 | 196 | Archaeal flagellin ArlB | 4 | 1 | 7-29 | 40-196 | 39762 (2) |
| MJ1_0707 | 632 | S-layer domain-like protein | 13 | 5 | 12-33, 598-617 | 29-598 | 6727 (12) |
| MJ1_0449 | 283 | Archaeal flagellin ArlB | 6 | 1 | 7-29 | 25-283 | 2700 (51) |
| MJ1_0408 | 476 | Hypothetical protein | 14 | 4 | 7-26, 326-348 | 28-325 | 1820 (107) |
| MJ1_0239 | 276 | Hypothetical protein | 5 | 2 | 7-29 | 25-276 | 1484 (141) |
| MJ1_0451 | 1132 | Pectin lyase-like protein | 48 | 6 | 12-34 | 30-1132 | 1479 (142) |
| MJ1_0467 | 328 | Hypothetical protein | 6 | 2 | 7-24 | 32-328 | 1413 (149) |
| MJ1_0701 | 908 | Pectin lyase-like protein | 36 | 1 | 7-29, 891-907 | 25-890 | 1358 (160) |
| MJ1_0179 | 418 | Lectin-like protein | 13 | 1 | 7-29 | 27-418 | 1263 (177) |
| MJ1_0691 | 1255 | Hypothetical protein | 27 | 7 | 16-27, 1238-1254 | 28-1237 | 1110 (217) |
| MJ1_0095 | 1410 | Hypothetical protein | 49 | 7 | 494-513, 518-537, 1393-1409 | 19-493, 538-1392 | 1046 (242) |
| MJ1_0625 | 94 | Hypothetical protein | 5 | 0 | 5-27 | 24-94 | 1037 (247) |
| MJ1_0420 | 161 | Archaeal flagellin-like protein | 4 | 1 | 7-29 | 24-161 | 985 (254) |
| MJ1_0153 | 228 | Metal-dependent hydrolase | 1 | 1 | N.D. | 20-228 | 869 (310) |
| MJ1_0375 | 403 | Glycosidases | 13 | 1 | 7-24 | 25-403 | 762 (367) |
| MJ1_0361 | 770 | Thermopsin | 26 | 6 | 7-29, 749-768 | 22-749 | 690 (418) |
| MJ1_0074 | 41 | Hypothetical protein | 0 | 0 | N.D. | 20-41 | 657 (442) |
| MJ1_0076 | 775 | Thermopsin | 42 | 5 | 754-774 | 21-753 | 656 (446) |
| MJ1_0309 | 479 | S-layer domain-like protein | 17 | 4 | 452-471 | 20-451 | 651 (451) |
| MJ1_0466 | 284 | Hypothetical protein | 4 | 1 | 4-21 | 20-284 | 650 (455) |
| MJ1_0299 | 514 | S-layer domain-like protein | 17 | 4 | 7-29, 482-504 | 20-482 | 641 (464) |
| MJ1_0108 | 285 | Hypothetical protein | 11 | 0 | 4-23, 263-284 | 24-262 | 629 (477) |
| MJ1_0435 | 211 | Hypothetical protein | 8 | 0 | 2-24 | 20-211 | 570 (527) |
| MJ1_0776 | 1320 | Pectin lyase-like protein | 52 | 8 | 7-24 | 27-1320 | 498 (577) |
| MJ1_0757 | 92 | Hypothetical protein | 4 | 0 | 5-22 | 22-92 | 332 (668) |

The predicted 3D protein structures for some of the putative cell surface proteins (Fig. 3) illuminate the structural characteristics, such as Ig (or PKD)-like fold consisting of a 2-layer sandwich of 7-9 antiparallel β-strands (52, 53) and pectin lyase-like folds consisting of parallel β strands coiled into a large helix (54), for each protein. It should be noted that the presence of these structural characteristics could not be addressed by the sequence homology-based analyses.

**Fig. 3.**
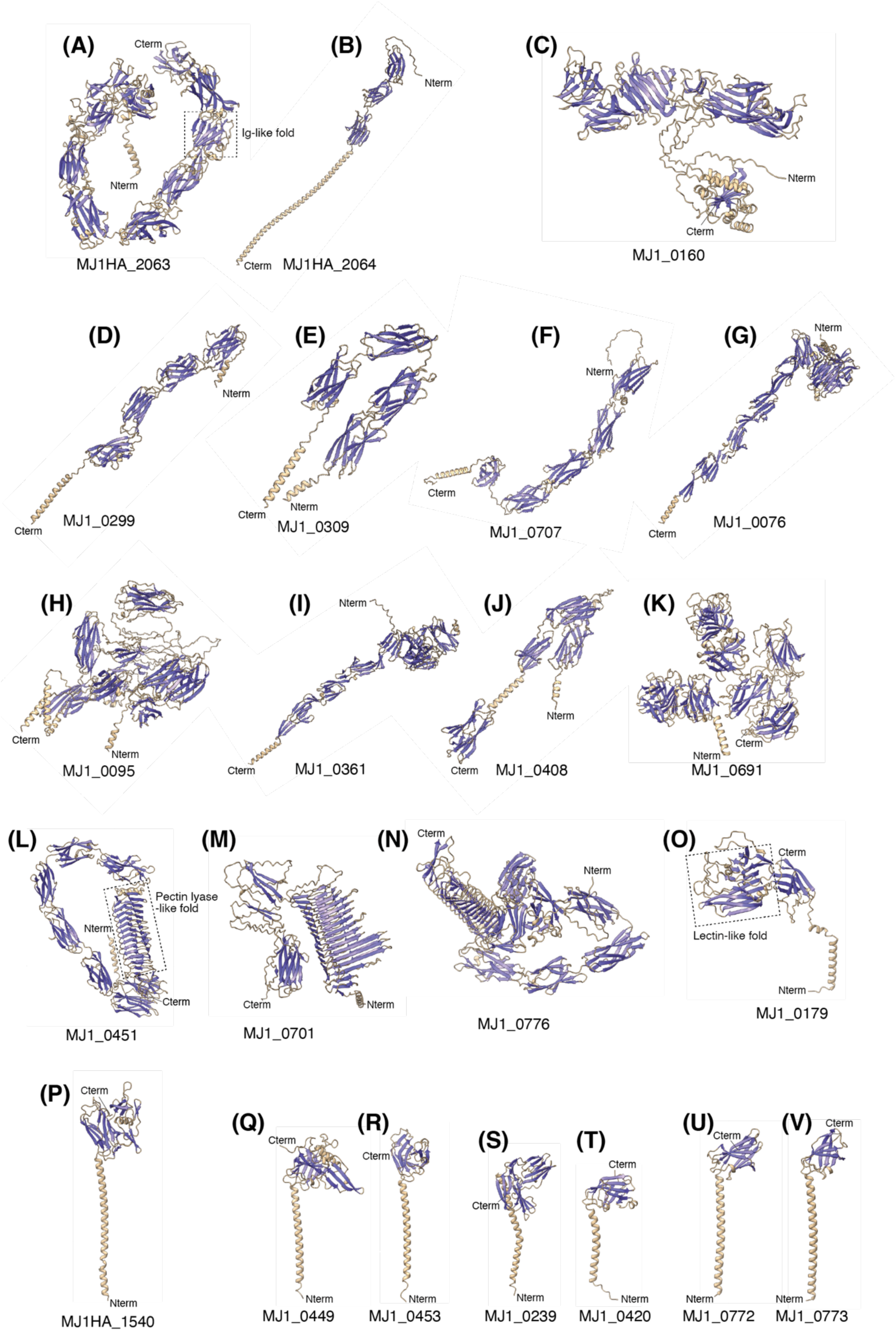
Predicted 3D structures of putative cell surface proteins. **(A)** SlaA and (B) SlaB of S-layer proteins of *Metallosphaera sedula* strain MJ1HA. The typical 3D structure of Ig-like fold is indicated in the box in the panel A. **(C–O)** Putative cell surface proteins with multiple glycosylation sites and Ig-like folds of *Nanobdella aerobiophila* strain MJ1^T^. The typical structures of pectin lyase- and lectin-like fold are indicated in the boxes in the panels L and O. **(P)** Archaellum component protein ArlB of strain MJ1HA, and **(Q–V)** ArlB-like proteins of strain MJ1^T^.

### S-layer proteins

For the host archaeon strain MJ1HA, the amino acid sequences from two genes (MJ1HA_2063 and MJ1HA_2064) were mostly identical to those of the S-layer proteins of SlaA and SlaB (99.2% and 100% amino acid sequence identities, respectively) of *Metallosphaera sedula* strain TH2^T^ (55). The predicted 3D protein structures of the SlaA and SlaB homologs of the strain MJ1HA (Fig. 3A and 3B) were also similar to the protein structures of SlaA and SlaB of *Sulfolobus acidocaldarius* that are assembly and anchoring domains of S-layers, respectively, as determined by cryo-EM (56, 57). The two genes were ranked in the top 5 of the highest expressed genes both in pure culture and in coculture with the strain MJ1^T^ (Table 1). The predicted 3D protein structures of the MJ1HA SlaA and SlaB homologs were characterized by many Ig-like folds and glycosylation sites, as consistent with other archaeal S-layer proteins including *S. acidocaldarius* SlaA and SlaB (56–59).

In contrast, no sequence homolog of any identified nor predicted S-layer proteins was found in the MJ1^T^ genome. This is not surprising, given that amino acid sequence identities of S-layer proteins are generally low among archaea in genus or higher taxonomic levels (60). No S-layer proteins have been determined for the DPANN archaea, although several candidates have been proposed (12, 13, 19, 20). We surveyed putative cell-wall or extracellular proteins with signal peptides (Table 1), and found many N-glycosylation sites, but no O-glycosylation sites, in the putative cell surface or extracellular proteins from the MJ1^T^ genes as described below.

A gene (MJ1_0160) annotated as a hypothetical protein was most highly expressed in the MJ1^T^ mRNA transcriptome (Table 1). The MJ1_0160 protein (1000 amino acids [a.a.]) was predicted to represent 15 glycosylation sites, 5 Ig-like folds (Fig. 3C), and a long extracellular region (>950 a.a.). No C-terminal TM helix was identified. These features suggest that MJ1_0160 encodes the S-layer assembly protein, like SlaA. Its homologs were only hit in several nanoarchaeal and other DPANN genomes by the BLASTp homology search (Table S3). Indeed, the MJ1_0160 protein showed a low similarity to the reported putative S-layer proteins of ‘*Nanoarchaeum equitans’* (NEQ300; 21.3% identity, e-value of 1e-10) and *‘Micrarchaeum harzensis’* (Micr_00292; 25.5% identity, e-value of 0.038). The predicted 3D protein structures of the NEQ300 and Micr_00292, which have been available in UniProt (61), represent multiple Ig-like folds (Fig. S7A). In addition, these genes have been highly expressed in mRNA- and protein levels, and predicted to represent multiple N-glycosylation sites (19, 22).

In addition, several genes potentially encoding S-layer-like proteins with multiple glycosylation sites and Ig-like folds were found in the MJ1^T^ genome (Table 1). For instance, three genes (MJ1_0299, MJ1_0309, and MJ1_0707) were predicted to represent S-layer domain proteins with 13–17 N-glycosylation sites, 4–5 Ig-like folds, and a C-terminal transmembrane (TM) helix (Fig. 3D–F). The N-terminal TM helix for the MJ1_0299 and MJ1_0707 proteins might be included in signal peptides that could be cleaved by a signal peptide (MJ1_0412; Table S1) in the maturation process. In particular, MJ1_0707 was highly expressed (ranked 12th), implying that it encodes a major S-layer anchoring protein, like SlaB.

### Other cell surface proteins

Five genes (MJ1_0076, MJ1_0095, MJ1_0361, MJ1_0408, and MJ1_0691) annotated as thermopsin or hypothetical proteins were predicted to represent 14–49 N-glycosylation sites, 4–7 Ig-like folds, and a C-terminal TM helix (Fig. 3G–K), which were not very highly expressed (ranked below 100th). Given that thermopsin is a peptidase (62), it is tempted to speculate that the thermopsin-like proteins from MJ1_0076 and MJ1_0361 are involved in the degradation of host S-layer proteins for constructing the symbiotic relationship, and are related to the observed unique structure between strains MJ1^T^ and MJ1HA as described above (Fig. 1D).

Notably, we found three genes (MJ1_0451, MJ1_0701, and MJ1_0776) annotated as pectin lyase-like proteins, of which lengths were 908–1320 a.a. These pectin lyase-like proteins were predicted to represent 36–52 N-glycosylation sites, 1–8 Ig-like folds, and a pectin lyase fold consisting of parallel beta-helix repeats (Fig. 3L–N). Although little is known about roles of pectin lyase in archaea, considering the function of pectin lyase degrading polysaccharides of plant cell walls (63), these pectin lyase-like MJ1 proteins are potentially involved in the degradation of the glycosylated host cell wall for symbiosis, and are also related to the observed unique structure between MJ1 and MJ1HA (Fig. 1D).

We detected a gene (MJ1_0179) for lectin-like protein by sequence homology-based analysis, which was predicted to represent a sandwich structure of over 10 β-strands in two sheets (i.e., Legume or concanavalin A-like lectin domain), an Ig-like fold, and an N-terminal TM helix (Fig. 3O). In addition, such lectin domains were also found in the predicted 3D structures of the above putative S-layer proteins, i.e., MJ1_0160 (Fig. 3C), MJ1_0076 (Fig. 3G), and MJ1_0691 (Fig. 3K), although these domains could not be detected by sequence homology-based analysis. Indeed, four genes (LC1Nh_0399–401, 0423) for lectin-like proteins have been reported from the cultivated nanohaloarchaeon *‘Nanohalobium constans’* (13), of which the predicted 3D structures (Fig. S7B) have been available in UniProt (61). Given the role of lectins in bacterial pathogenic adherence (64), these lectin-like proteins are potentially involved in the recognition of and attachment to their hosts, as mentioned previously (13).

### Archaellum-related proteins

In addition to S-layer proteins, genes for archaellum-related Arl (formerly Fla) proteins (65), i.e., ArlBFGHIJX (MJ1HA_1534 to 1540), were found in the genome of the host archaeon strain MJ1HA (Table S2). The gene (MJ1HA_1540) encoding ArlB was highly expressed in pure culture and coculture (ranked in the top 5). The predicted 3D structure of the putative ArlB represented an Ig-like fold, a long N-terminal α-helix (Fig. 3P), and several N-glycosylation sites, of which feature is a hallmark of archaellin, i.e., an extracellular filamentous part of the archaellum protein complex (66–69).

Genes for ArlBHIJ have been found in the MJ1 genome (Table S1) as reported previously (5). Of the two genes for ArlB homologs (MJ1_0449 and MJ1_0453), MJ1_0453 was the second highest expressed in the MJ1 transcriptome (Table 1), suggesting that it is a major archaellin protein. The predicted 3D structures of the two ArlB homologs showed the presence of an Ig-like fold and an N-terminal TM helix (Fig. 3Q and R), like ArlB of its host archaeon (MJ1HA_1540) (Fig. 3P). In addition, based on the 3D protein structures, additional two genes (MJ1_0239 and MJ1_0420) could be related to archaellum-like filaments (Fig. 3S and T). Moreover, the proteins for MJ1_0772 and MJ1_0773 were also predicted to represent an archaellin-like structure (Fig. 3U and V), although no signal peptides were detected. The detection of a variety of the genes for the archaellin-like proteins implies the presence of additional filaments of strain MJ1^T^ besides the typical archaella, like other archaea such as *Sulfolobus acidocaldarius* (26, 70) and *Pyrobaculum calidifontis* (25, 71). Indeed, we observed two types of extracellular filaments with different thicknesses by SEM and TEM (Fig. 2D and E)(5).

### Live imaging of cell motility

The host archaeon strain MJ1HA was weakly motile even at room temperature, as consistent with the presence of archaella (5). In this study, the live imaging of the MJ1HA pure culture indicated that MJ1HA cells were highly motile at 65°C with a swimming speed of 8.0 ± 1.9 μm/s (N = 35 cells)(Movie S4). The live imaging of the MJ1–MJ1HA coculture indicated that MJ1–MJ1HA cell complexes were also motile with 13.0 ± 4.2 µm/s (N = 11 cells)(Movie S5). In contrast, sole MJ1 cells without host attachment were not motile under the conditions, despite the presence of archaellum-like filaments revealed by TEM (Fig. 2E; Fig. S6). Strain MJ1^T^ may use the filaments for attachment to their host cells that are swimming and occasionally encounter MJ1^T^ cells, but not for swimming which is a highly-energy-consuming activity. This is reasonable because strain MJ1^T^ cannot generate ATP by itself as indicated by the genome analysis, which depends on the uptake from its host, and must serve energy for survival.

### A hypothetical model of cell surfaces and symbiosis

Based on the results of the microscopic and molecular analyses, we propose a hypothetical model for the cell surface architecture and symbiotic lifestyle of *Nanobdella aerobiophila* strain MJ1^T^ and its host archaeon *Metallosphaera sedula* strain MJ1HA (Fig. 4). The cells of strain MJ1HA are probably covered with the highly expressed SlaA and SlaB homologs as the major S-layer proteins (Fig. 4A), like *Metallosphaera sedula* strain TH2^T^ and other relatives in the order *Sulfolobales* (55, 56). Typical archaellin, i.e., ArlB, are also present on its surface. The proteins of the S-layer and archaellin are likely to be highly glycosylated, which could serve as the targets for attachment by strain MJ1^T^.

**Fig. 4.**
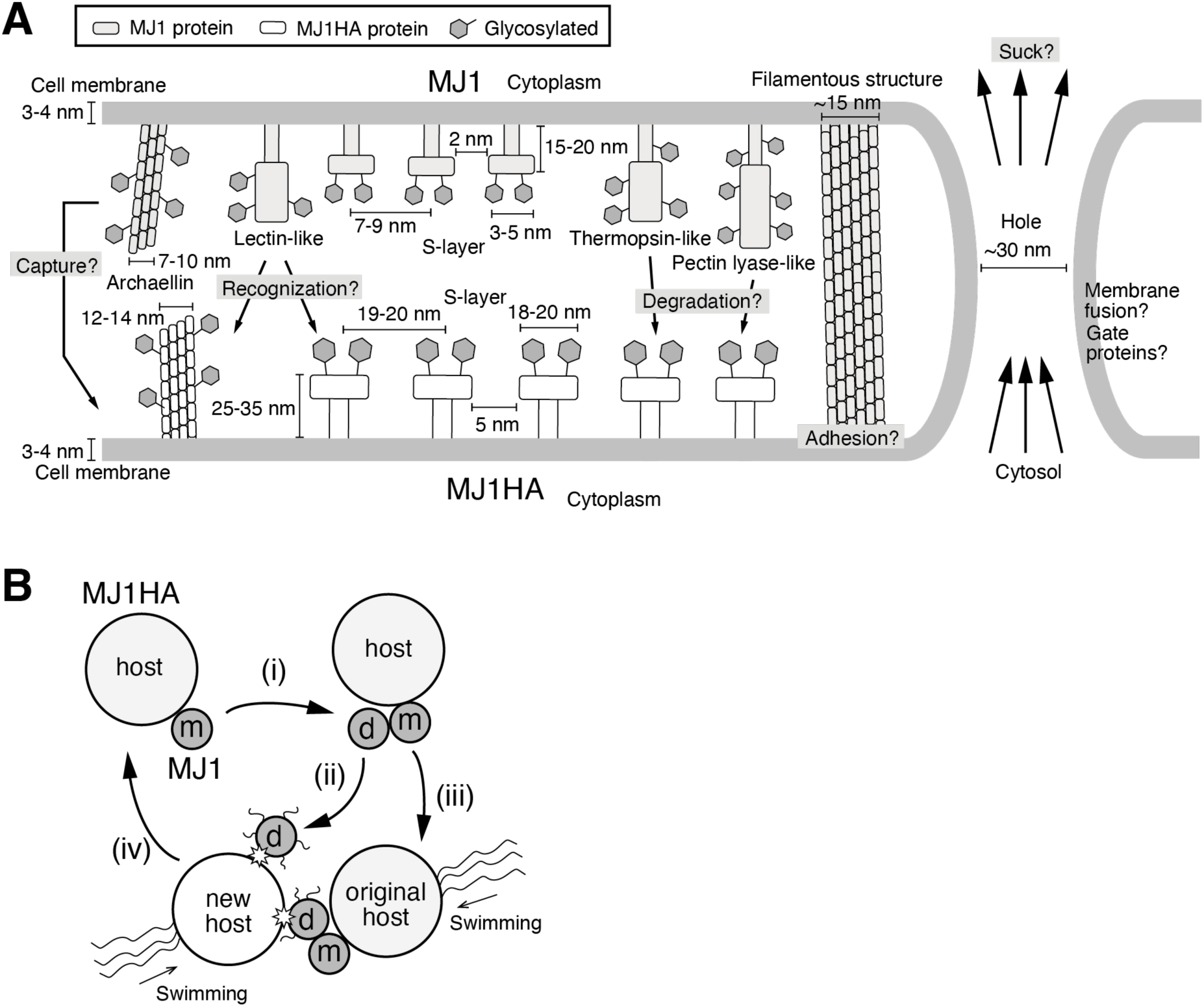
Proposed model for the cell surface architecture and symbiotic lifestyle of *Nanobdella aerobiophila* strain MJ1^T^ and its host archaeon *Metallosphaera sedula* strain MJ1HA. (**A**) Putative cell surface proteins involved in cell-cell interaction between strains MJ1^T^ and MJ1HA. **(B)** Proposed life cycle of strain MJ1^T^. m, mother cells. d, daughter cells. (i) A MJ1^T^ cell is divided on the cell surface of its host. The daughter cell (ii) leaves the host or (iii) swims with the host, attaches to a new host cell, and (iv) sucks the cytosol from the host.

The S-layer lattice symmetry and constant of strain MJ1^T^ are different from those of its host archaeon MJ1HA and even the other reported DPANN archaea (18, 19). Strain MJ1^T^ possesses no obvious homologs to known S-layer proteins. Our structure-based prediction and transcriptome analyses suggest that the MJ1_0160 protein serves as the major S-layer assembly protein like SlaA. The MJ1_0707 protein (or other candidates such as MJ1_0229 and MJ1_309) may link between the assembly proteins and cell membrane as the S-layer anchoring protein like SlaB. Archaellum-like filaments consisting of ArlB are also present on its surface. Additional cell surface proteins are likely to be present, which include pectin lyase-, thermopsin-, and lectin-like proteins. These proteins are potentially related to the observed irregular structures (larger globules, flat sheets, and fibers) by QFDE-TEM besides the S-layer and archaella, which could be involved in the recognition and attachment to the host, and/or degradation of the host S-layer. In particular, the lectin-like MJ1_0179 protein could be used for the recognition of the specific host; strain MJ1^T^ constructs a symbiotic relationship only with strain MJ1HA, but not others (5), like the nanoarchaeon *‘Nanoarchaeum equitans’* capable of growing only with its host *Ignicoccus hospitalis* (4). Although MJ1^T^ cells are equipped with archaellum-like filaments, they cannot swim alone, and thus wait to be hit by the swimming cells of its host archaeon strain MJ1HA (Fig. 4B). After attachment and sucking the cytosol from the host, MJ1^T^ prolificate on the cell surface of its host, and may leave for free. This lifestyle of MJ1^T^ (and maybe some other DPANN archaea), if it is true, resembles that of phages or viruses.

The proposed model is reasonable, although still speculative. Further experimental evaluation (e.g., protein characterization, gene manipulation, and live imaging of material transfer from a host to a symbiont) is needed to reconstruct a more reliable model. In addition, there are some limitations to the proposed model. In this study, we focused only on putative cell surface proteins with detectable signal peptides, and therefore, the proposed model could not include cell surface proteins with unrecognized signal peptides or without signal peptides, even if such cell surface proteins were present. Besides, it is still largely unclear how a MJ1^T^ cell constructs the hole penetrating the cell membranes of MJ1^T^ and MJ1HA cells, and how it sucks the host cytosol if it is true. These mechanisms might be unique among archaea (and even all living things) and will open a new window into the research fields of microbial physiology and ecology. Strain MJ1^T^ capable of well growing under aerobic conditions is easy to cultivate, and will contribute to elucidating the symbiotic mechanisms as a model organism for DPANN archaea.

## Acknowledgments

We are grateful to Ms. Nagisa Sato and Mr. Minoru Fukushima for their technical support on RNA-seq and Tukey Window processing, respectively. This work was supported by the Institute of Fermentation (IFO), JSPS KAKENHI Grant Number 19H05679, 19H05688, 19H05689 (Post-Koch Ecology), 22K19141, 22H04881, JST CREST JPMJCR19S2, and the RIKEN interdisciplinary research program Integrated Symbiology (iSYM).

## Data availability

The raw data for RNA-seq was deposited in DDBJ under the accession number DRA011278.

